# A critical role for Musashi in photoreceptor morphogenesis and Vision

**DOI:** 10.1101/2020.01.25.919761

**Authors:** Jesse Sundar, Fatimah Matalkah, Bohye Jeong, Peter Stoilov, Visvanathan Ramamurthy

## Abstract

Musashi family of RNA-binding proteins are known for their role in stem-cell renewal and are negative regulators of cell differentiation. Interestingly, in the retina, Musashi proteins, MSI1 and MSI2 are differentially expressed throughout the cycle of retinal development including robust expression in the adult retinal tissue. To study the role of Musashi proteins in the retina, we generated a pan-retinal and rod photoreceptor neuron specific conditional knockout mouse lacking MSI1 and MSI2. Independent of sex, photoreceptor neurons with simultaneous deletion of *Msi1* and *Msi2* were unable to respond to light and displayed severely disrupted OS morphology and ciliary defects. The retina lacking Musashi exhibited neuronal degeneration with complete loss of photoreceptors by six months. In concordance with our earlier studies that proposed a role for Musashi in regulating alternative splicing, the loss of Musashi prevented the use of photoreceptor-specific exons in transcripts critical for OS morphogenesis, ciliogenesis and synaptic transmission. Overall, we demonstrate a critical role for Musashi in the morphogenesis of terminally differentiated photoreceptor neurons. This role is in stark contrast with the canonical function of these two proteins in maintenance and renewal of stem cells.

## INTRODUCTION

In eukaryotes, alternative splicing of pre-mRNA increases protein diversity and controls gene expression. Diversification of proteomes through alternative splicing is a defining characteristic of metazoans and was expanded dramatically in bilaterians (1). Alternative splicing is prevalent in vertebrate neurons and is critical for the development and function of vertebrate nervous systems (2–7).

We previously showed that photoreceptor neurons exploit a unique splicing program (8). Motif enrichment analysis suggested that Musashi-1 (MSI1) and Musashi-2 (MSI2) promote the use of photoreceptor specific exons (8). We further showed that MSI1 is critical for utilization of photoreceptor specific exon in the *Ttc8* gene (8). In addition, Musashi promotes the splicing of several photoreceptor specific exons when over-expressed in cultured cells (8). Recently, analysis of a comprehensive gene expression data set demonstrated that photoreceptors utilize a unique set of alternative exons that are primarily regulated by MSI1 and MSI2 (9).

The MSI1 and MSI2 proteins have two highly conserved RNA binding domains (RBDs) in the N-terminal region which show close to 90% sequence identity and recognize a similar UAG motif in RNA (10). The two RBDs of MSI1 and MSI2 are followed by a less conserved C-terminal region which shows approximately 70% sequence identity (11). The high degree of sequence identity between the MSI1 and MSI2 results in functional redundancy between the two proteins (12, 13).

Vertebrate photoreceptors are neurons specialized in detecting and transducing light stimuli. Photoreceptors are characterized by segmented morphology which compartmentalizes phototransduction, core cellular functions, and synaptic transmission. The light sensing machinery is confined to the outer segment, a stack of membranes that is elaborated by cell’s modified primary cilium. The outer segment is dynamic structure that is remade every 7 to 10 days. Consequently, maintenance of the outer segment requires high rate of transport of membranes and proteins through the connecting cilium (14).

The predicted splicing targets of Musashi in photoreceptors include pre-mRNAs from ciliary (*Ttc8, Cep290, Cc2d2a, Prom1*) and synaptic-associated genes (*Cacna2d4, Slc17a7*) (15–21). These genes are crucial for photoreceptor development and function (15–21). We proposed that production of photoreceptor specific splicing isoforms that is promoted by Musashi is necessary for the development and maintenance of photoreceptor cells *in vivo* (8).

To test if Musashi drives photoreceptor development and function, we removed *Msi1* and *Msi2* in the developing retina and rod photoreceptor cells. We find that Musashi proteins are essential for photoreceptor function, morphogenesis, and survival but not their specification. Specifically, the Musashi proteins are crucial for outer segment (OS) and axoneme development. As expected, disruption of the Musashi genes led to loss of expression of photoreceptor specific splicing isoforms.

## MATERIALS AND METHODS

### Generation of mice and genotyping

Mice carrying floxed alleles for *Msi1* and *Msi2* were provided by Dr. Christopher Lengner from the University of Pennsylvania. *Six3-Cre* transgene or *Nrl-Cre* transgenes were used to delete the floxed alleles in the developing retina or rod photoreceptors (Stock Nos. 019755, 028941, Jax labs). All mouse lines in this study are in C57 Black6/J background (https://www.jax.org/strain/000664) and were devoid of naturally occurring *rd1* and *rd8* alleles (22, 23). Males hemizygous for the *Six3-Cre* transgene or *Nrl-Cre* transgene and floxed for either *Msi1*, *Msi2*, or both *Msi1* and *Msi2* were mated with females floxed for either *Msi1*, *Msi2*, or both *Msi1* and *Msi2* to obtain experimental knockout mice and littermate control. The offspring of breeding pairs were genotyped using PCR from ear biopsies. The *Msi1* wildtype and floxed alleles were identified using following primers: (5’-CGG ACT GGG AGA GGT TTC TT-3’ and 5’-AGC TCC CCT GAT TCC TGG T-3’). The *Msi2* wildtype and floxed alleles were identified by using following primers: (5’-GCT CGG CTG ACA AAG AAA GT-3’ and 5’-TCT CCT TGT TGC GCT CAG TA-3’). The presence of the *Six3 Cre, Nrl Cre* transgene and *Cre recombinase* were determined using following primers respectively: (5’-CCC AAA TGT TGC TGG ATA GT-3’ and 5’-CCC TCT CCT CTC CCT CCT-3’), (5’-TTT CAC TGG CTT CTG AGT CC-3’ and 5’-CTT CAG GTT CTG CGG GAA AC-3’) and (5’-CCT GGA AAA TGC TTC TGT CCG-3’ and 5’-CAG GGT GTT ATA AGC AAT CCC-3’).

All experiments were conducted with the approval of the Institutional Animal Care and Use Committee at West Virginia University. All experiments were carried out with adherence to the principles set forth in the ARVO Statement for the Ethical Use of Animals in Ophthalmic and Vision Research which advocates the use of the minimum number of animals per study needed to obtain statistical significance.

### Electroretinography, Immunoblotting, and Reverse Transcriptase PCR

Electroretinography, immunoblotting, and reverse transcriptase PCR were conducted using previously described protocol from our laboratory (8, 24, 25).

### Immunofluorescence Microscopy

Immunofluorescence microscopy was carried out using a modified procedure in our laboratory(24, 25). Briefly, eyes were enucleated, and the cornea and lens were discarded. After dissection, eyes were fixed by immersion in 4% paraformaldehyde in PBS for one hour. After washing the eyes in PBS three times for ten minutes each, they were dehydrated by overnight incubation in 30% sucrose in PBS. Eyes were then incubated in a 1:1 solution of OCT:30% sucrose in PBS for one hour and frozen in OCT (VWR). The frozen tissues were sectioned using a Leica CM1850 cryostat for collecting serial retinal sections of 16μm thickness. The retinal cross-sections were then mounted onto Superfrost Plus microscope slides (Fisher Scientific). Slide sections were then washed and permeabilized with PBS supplemented with 0.1% Triton X-100 (PBST) and incubated for one hour in a blocking buffer containing 10% goat serum, 0.3% Triton X-100, and 0.02% sodium azide in PBS. Retinal sections were then incubated with primary antibody in a dilution buffer containing 5% goat serum, 0.3% Triton X-100, 0.02% sodium azide, and primary antibody at 1:500 dilution in PBS overnight at 4°C followed by three 5-minute washes using PBST. Sections were then incubated in the same dilution buffer containing secondary antibody and DAPI at 1:1000 for one hour. Slides were washed with PBST three times for five minutes each before treating with Prolong Gold Antifade reagent (ThermoFisher) and securing the coverslip. The images were collected using a Nikon C2 Confocal Microscope.

### Retinal histology of the mouse models

Following euthanasia, eyes were enucleated using C-shaped forceps after marking the superior pole and incubated in Z-fixative for >48 hours before shipment and tissue processing by Excalibur Pathology Inc. (Norman, OK). The embedding, serial sectioning, mounting, and hematoxylin/eosin (H&E) staining were performed by Excalibur Pathology. A Nikon C2 Microscope equipped with Elements software was used to image the slides.

### Transmission Electron Microscopy

After euthanasia, a C-shaped forceps was used to enucleate the eye, and the cornea was discarded (24, 25). Eyes were then incubated in a fixative solution containing 2.5% glutaraldehyde and 2% paraformaldehyde in 100mM sodium cacodylate buffer at pH 7.5 for 45 minutes before removal of the lens. After lensectomy, eyes were placed back into fixative for 72 hours before shipment, tissue processing, and imaging at the Robert P. Apkarian Integrated Electron Microscopy Core at Emory University.

### Antibodies and stains

The following primary antibodies were used throughout our studies: rat anti-MSI1 (1:1000; MBL International Cat# D270-3, RRID:AB_1953023), rabbit anti-MSI2 (1:2000; Abcam Cat# ab76148, RRID:AB_1523981), mouse anti-α-tubulin (1:10 000; Sigma-Aldrich Cat# T8328, RRID:AB_1844090), rhodamine peanut agglutinin (1:1000; Vector Laboratories Cat# RL-1072, RRID:AB_2336642), rabbit anti-peripherin-2 (1:2000) was a kind gift by Dr. Andrew Goldberg from Oakland University, rabbit anti-PDE6β (1:2000; Thermo Fisher Scientific Cat# PA1-722, RRID:AB_2161443), mouse anti-acetylated α-tubulin (1:1000; Santa Cruz Biotechnology Cat# sc-23950, RRID:AB_628409), guinea pig anti-MAK (1:500; Wako, Cat# 012-26441, RRID:AB_2827389), mouse anti-glutamylated tubulin (1:500; AdipoGen Cat# AG-20B-0020B, RRID:AB_2490211), mouse anti-Ttc8 (1:1000; Santa Cruz Biotechnology Cat# sc-271009, RRID:AB_10609492), rabbit anti-Ttc8 Exon 2A (1:1000; Peter Stoilov, West Virginia University, Cat# Anti-Bbs8 exon 2A, RRID:AB_2827390), mouse anti-GAPDH (1:10,000; Fitzgerald Industries International Cat# 10R-G109a, RRID:AB_1285808), and 4′,6-diamidino-2-phenylindole (DAPI: nuclear counterstain; 1:1000; ThermoFisher, Waltham, MA).

### Statistical analysis

Unless otherwise stated, the data is presented as mean of at least three biological replicates with error bars representing the standard error of the mean. Statistical significance was determined by homoscedastic, two-tailed unpaired T-test.

## RESULTS

### Validation of the conditional knockout mouse models

We analyzed the expression of Musashi proteins in various tissues from adult mice. Out of all the tissues we tested, retina showed the highest expression of MSI1 and MSI2 proteins (Figure 1A), in line with the previously reported high transcript levels for *Msi1* and *Msi2* in rod photoreceptors (9). To test the biological significance of Musashi protein expression in the murine retina, we used *Cre-LoxP* conditional recombination to remove either *Msi1*, *Msi2*, or both the *Msi1* and *Msi2* genes throughout the entire retina and ventral forebrain using the *Six3 Cre* transgene (26). Throughout this work, we refer to *Musashi* floxed mice which are hemizygous for the *Six3 Cre* transgene as *ret-Msi-/-* mice. The conditional recombination results in the deletion of *Msi1’s* transcription start site, exon 1, and exon 2 (13). For *Msi2*, the transcription start site and the first four exons are removed after cre-mediated recombination (13). The ablation of MSI1 and MSI2 was confirmed by immunoblotting retinal lysates from knockout mice at postnatal day 10 (PN10) (Figure 1B). Immunofluorescence microscopy of retinal cross sections obtained from the knockout mice affirmed the absence of MSI1 and MSI2 expression in the retina (Figure 1C).

**Figure 1:**
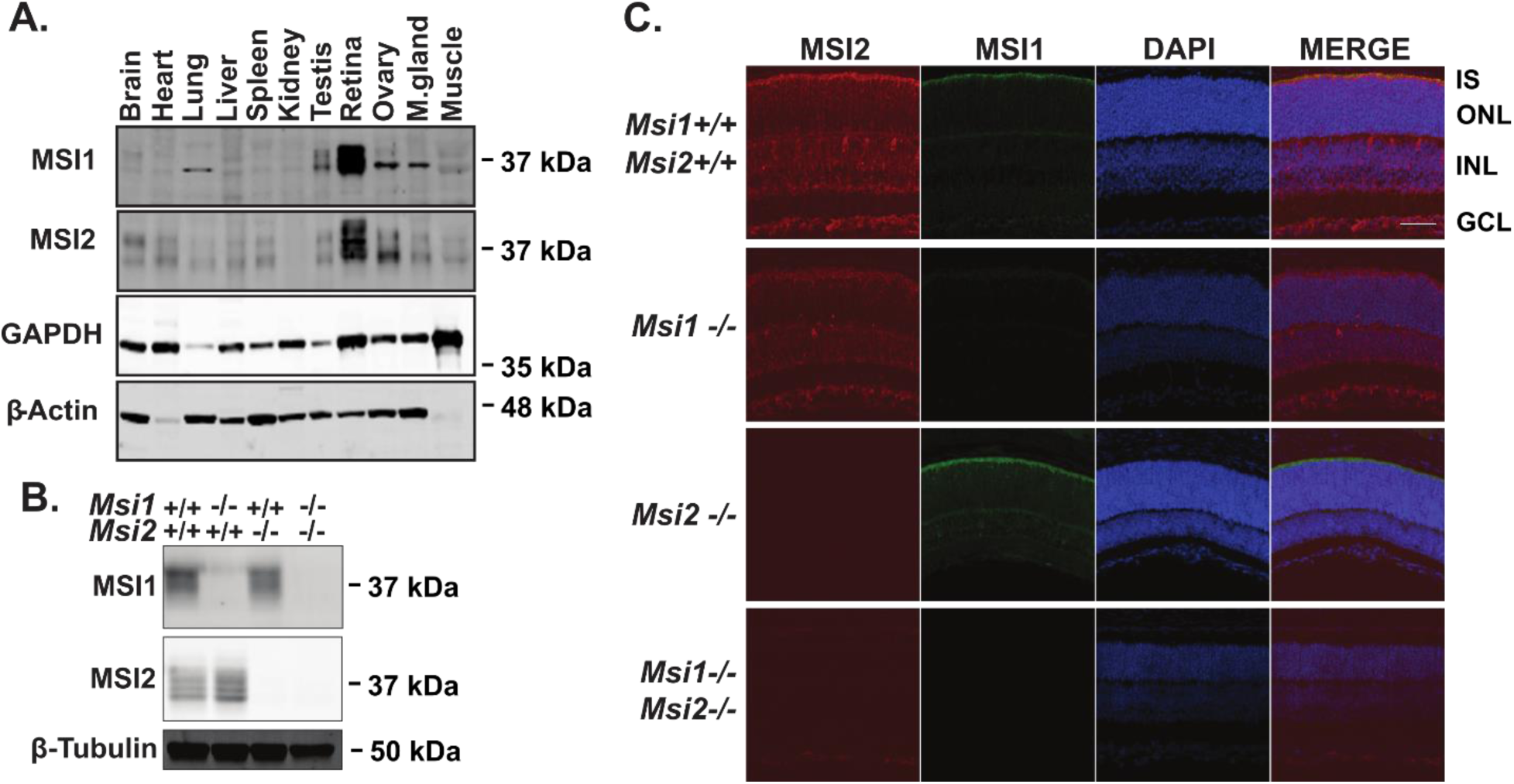
Conditional Musashi knockout mouse models. **A.**Immunoblot of indicated tissues from adult wildtype mice probed with MSI1 and MSI2 antibodies. GAPDH and β-Actin serve as a loading control. **B.**Western blot analyses of Musashi in retinal lysates from *ret-Msi1-/-*, *ret-Msi2-/-*, and *ret-Msi1-/-: Msi2-/-* mice at PN10. β-tubulin levels provide a loading control. **C.** Retinal sections from *ret-Msi1-/-*, *ret-Msi2-/-*, and *ret-Msi1-/-: Msi2-/-* mice at PN10 probed with MSI1 (Green) and MSI2 (Red) antibodies along with a DAPI nuclear counterstain (Blue). (IS: inner segment, ONL: outer nuclear layer, INL: inner nuclear layer, and GCL: ganglion cell layer). Scale bar = 50 µm.

### The Musashi proteins are crucial for photoreceptor function

To determine if the Musashi proteins are required for photoreceptor function, we performed electroretinographic (ERG) recordings of the *Musashi* conditional knockout mice at PN16 and monitored for changes in retinal function up to PN180. Figure 2A shows the representative scotopic and photopic ERG waveforms of the *ret-Msi1-/-*, *ret-Msi2-/-*, and *ret-Msi1-/-:Msi2-/-* mice at PN16 immediately after mice open their eyes (27). When both *Musashi* genes are removed, no scotopic or photopic response remains as shown by absence of conspicuous “a”-waves and “b”-waves (Figure 2A). However, significant photoreceptor function remains in the *ret-Msi1-/-* and *ret-Msi2-/-* single knockout mice. We characterized the photoreceptor function of the *ret-Msi1-/-* and *ret-Msi2-/-* mice further to see if there was a progressive loss of vision as the mice aged (Figure 2B–E). In *ret-Msi1-/-* mice, there was a statistically significant reduction in photoreceptor “a”-wave amplitudes at almost all light intensities (Figure 2B). This reduction in the photoreceptor “a”-wave amplitude was stationary and persisted in *ret-Msi1-/-* mice up to PN180 (Figure 2C). On the other hand, *ret-Msi2-/-* mice at PN16 had normal photoreceptor function at all the light intensities we tested (Figure 2D). The “a”-wave amplitude began to decrease progressively in *ret-Msi2-/-* mice as they aged, and this became significant at PN120 (Figure 2E).

**Figure 2:**
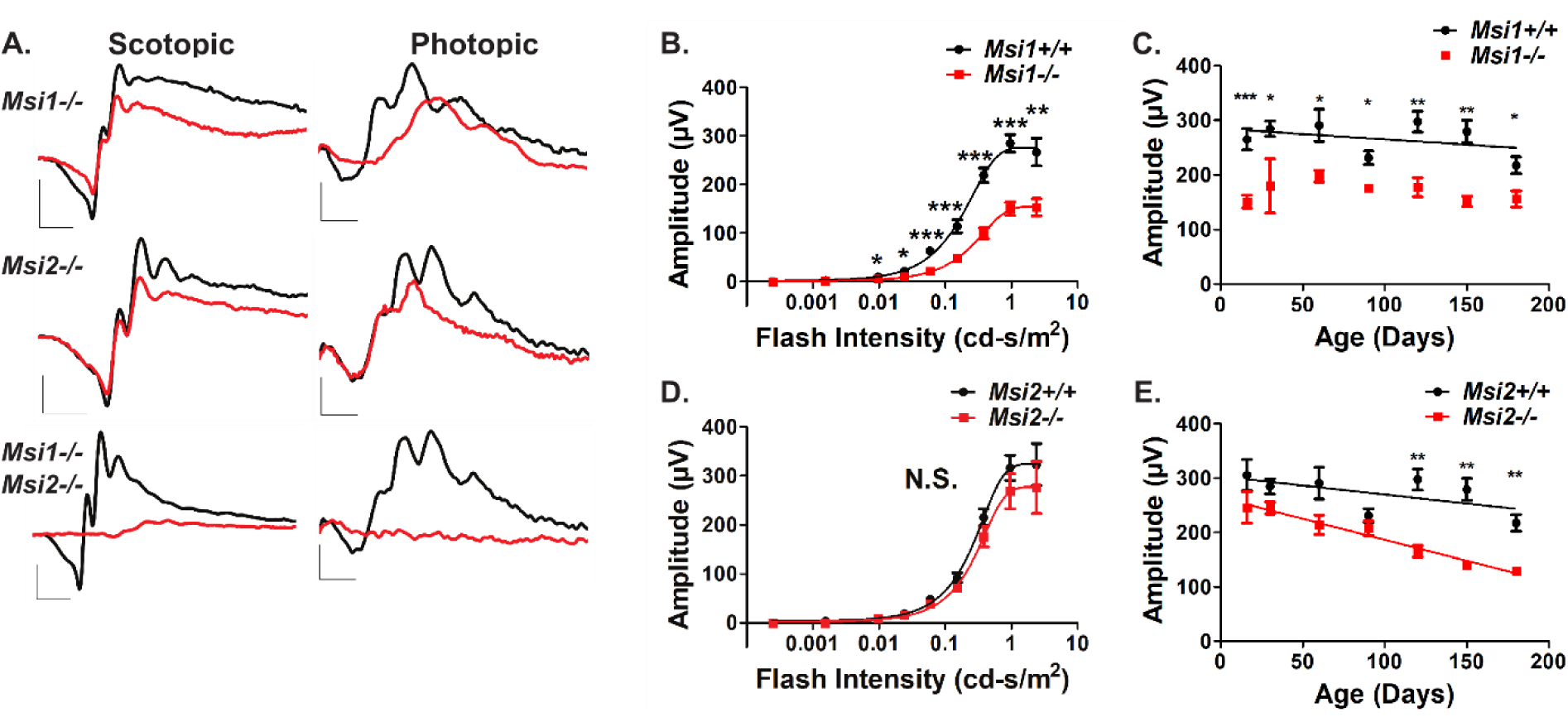
The Musashi proteins are crucial for normal visual response. **A.** Representative scotopic and photopic electroretinograms (ERGs) from the *ret-Msi1-/-*, *ret-Msi2-/-*, and *ret-Msi1-/-: Msi2-/-* mice at PN16. Scotopic ERGs were obtained after overnight dark adaptation using 0.151 cd-s/m^2^ flashes while photopic ERGs were obtained with 7.6 cd-s/m^2^ flashes under light-adapted conditions using a rod-saturating white background light (Scotopic scale bar: x-axis = 20ms, y-axis = 200 µV; Photopic scale bar: x-axis = 20ms, y-axis = 20 µV). **B.** Intensity-response plot of the scotopic “a”-wave response from *ret-Msi1-/-* mice (*=P-value < 0.05; **=P-value < 0.01; ***=P-value < 0.001). **C.** Plot of the rod photoreceptor “a”-wave response from *ret-Msi1-/-* mice against the age of the mouse during which the ERG was recorded. **D.** Intensity-response curve of the scotopic “a”-wave response from *ret-Msi2-/-* mice (*=P-value < 0.05; **=P-value < 0.01; ***=P-value < 0.001). **E.** Plot of the rod photoreceptor “a”-wave response from *ret-Msi2-/-* mice plotted against the age of the mouse during which the ERG was recorded. All data is shown as the mean ± the SEM, and statistical analyses were carried out using the homoscedastic unpaired student’s t-test (*=P-value<0.05).

The two Musashi protein share high degree of sequence similarity and are proposed to be functionally redundant, yet the progression of vision loss in the single *Msi1* and *Msi2* knockouts was significantly different. We tested if changes in expression levels of the two proteins after birth may account for this discrepancy. Western blot analysis of the Musashi protein expression levels in the retina between postnatal days 0 and 110, showed a distinct pattern of expression (Figure 3A and B). MSI1 levels spike by postnatal day 4 and remain high until P13-P16, a time frame that includes the period of photoreceptor outer segment morphogenesis (Figure 3A and B). After eye opening MSI1 protein expression declines (Figure 3A and B). MSI2 shows inverse pattern of protein expression to that of MSI1: relatively low levels after birth that gradually increase and peak after postnatal day 16 as the MSI1 protein levels decline (Figure 3A and B). Overall, our data shows that the Musashi proteins essential for photoreceptor function. The two proteins are functionally redundant, but appear to act at different time points of the retinal development.

**Figure 3:**
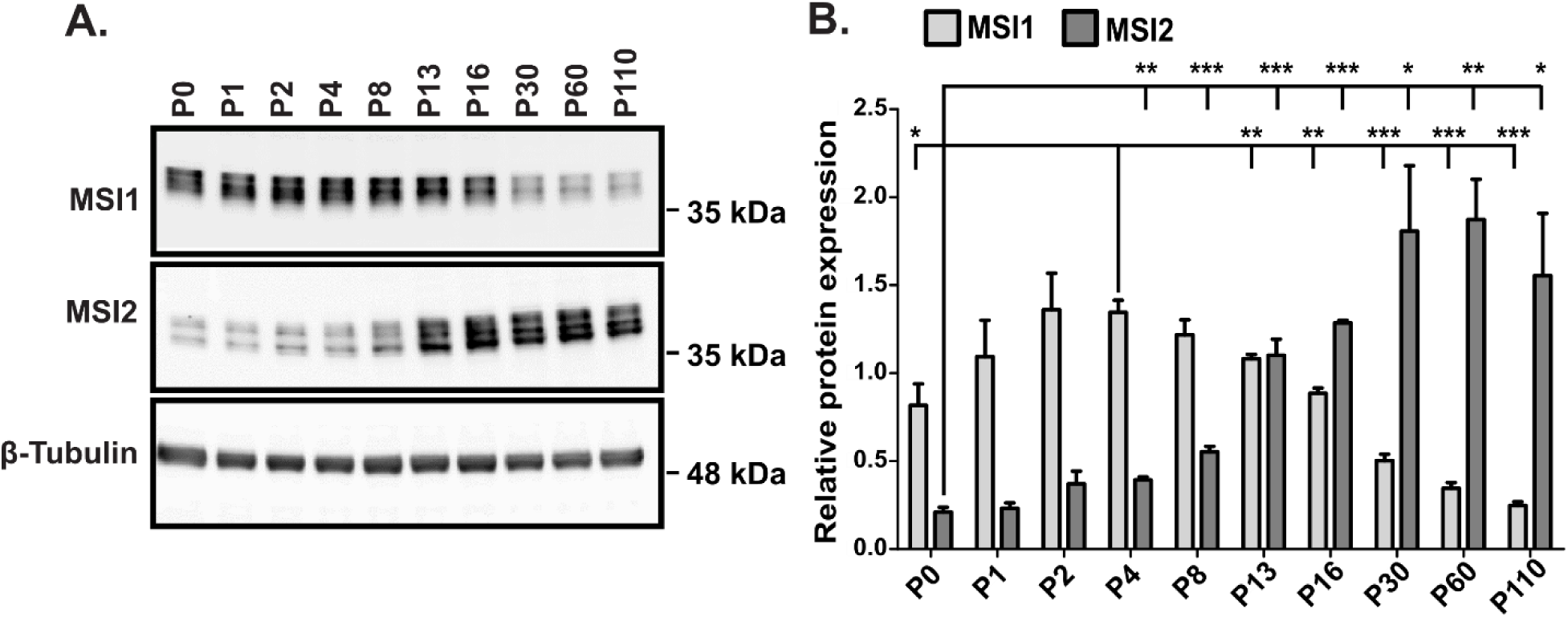
Developmental switch in expression of MSI 1 and 2. **A.** Representative immunoblot showing the expression of MSI1 and 2 in retinal tissues at indicated ages (P0-P110). Equal amount of total protein (20 µg) were loaded in each lane. β-tubulin serves as the loading control. **B.** Quantification of immunoblots shown in Panel A (n=3). All data is shown as the mean ± the SEM, and statistical analyses were carried out using the homoscedastic unpaired student’s t-test (*=P-value < 0.05; **=P-value < 0.01; ***=P-value < 0.001). T-test for Msi1 compared expression levels to the peak expression at P4. T-test for Msi2 compared expression levels to the expression at P0.

### Intrinsic expression of Musashi in photoreceptors is crucial for photoreceptor function

We next sought to determine if the phenotype of the *ret-Msi-/-* mice was due to the absence of Musashi protein expression in photoreceptors or if deletion of Musashi in other retinal cell types or retinal progenitors were contributing to the loss of vision. To this end, we generated rod-specific *Musashi* conditional knockouts by crossing *Musashi* floxed mice with mice hemizygous for the *Nrl Cre* transgene where the *Nrl* promoter activates Cre expression in rod photoreceptors (28). Throughout this work, we refer to the *Musashi* floxed mice that are hemizygous for the *Nrl Cre* transgene as *rod-Msi-/-* mice. We used ERG to analyze the retinal function of the knockout mice after ablation of the *Musashi* genes in rods (Figure 4 A–E). Figure 4A shows the scotopic and photopic ERG waveforms of the *rod-Msi1-/-*, *rod-Msi2-/*-, and *rod-Msi1-/-:Msi2-/-* mice at PN16. As observed in the *ret-Msi1-/-:Msi2-/-* mice, no significant rod function was observed in the *rod-Msi1-/-:Msi2-/-* mice at PN16 which is demonstrated by absence of conspicuous “a”-wave under scotopic testing conditions (Figure 4A). We examined the *rod-Msi1-/-* and *rod-Msi2-/-* single knockout mice to see if the photoresponse phenotype was comparable to that obtained from the *ret-Msi1-/-* and *ret-Msi2-/-* mice. In *rod-Msi1-/-* mice at PN16, there was a reduction in photoreceptor “a”-wave amplitudes at multiple light intensities (Figure 4B). This reduction in “a”-wave amplitude persisted as these mice aged up to PN180 (Figure 4C). Contrarily, PN16 *rod-Msi2-/-* mice had no changes in photoreceptor function at all the light intensities examined (Figure 4D). As observed in the *ret-Msi2-/-* mice, the “a”-wave amplitude began to decrease progressively as these mice aged, and this decrease became statistically significant at PN90 (Figure 4E). The similar phenotypes of the *ret-Msi* and *rod-Msi* knockout mice shows that the intrinsic expression of Musashi proteins in photoreceptors is crucial for their function and that deletion of Musashi proteins in other cell types likely does not contribute significantly to the phenotype observed in the *ret-Msi-/-* mice. Therefore, throughout the rest of our studies, we focus on the *ret-Msi1-/-:Msi2-/-* mouse model for our experiments since there is a compensation in function occurring between MSI1 and MSI2 in the single knockout mice and to avoid confounding results that might be obtained when *Msi1* and *Msi2* are deleted only in rod but not cone photoreceptors.

**Figure 4:**
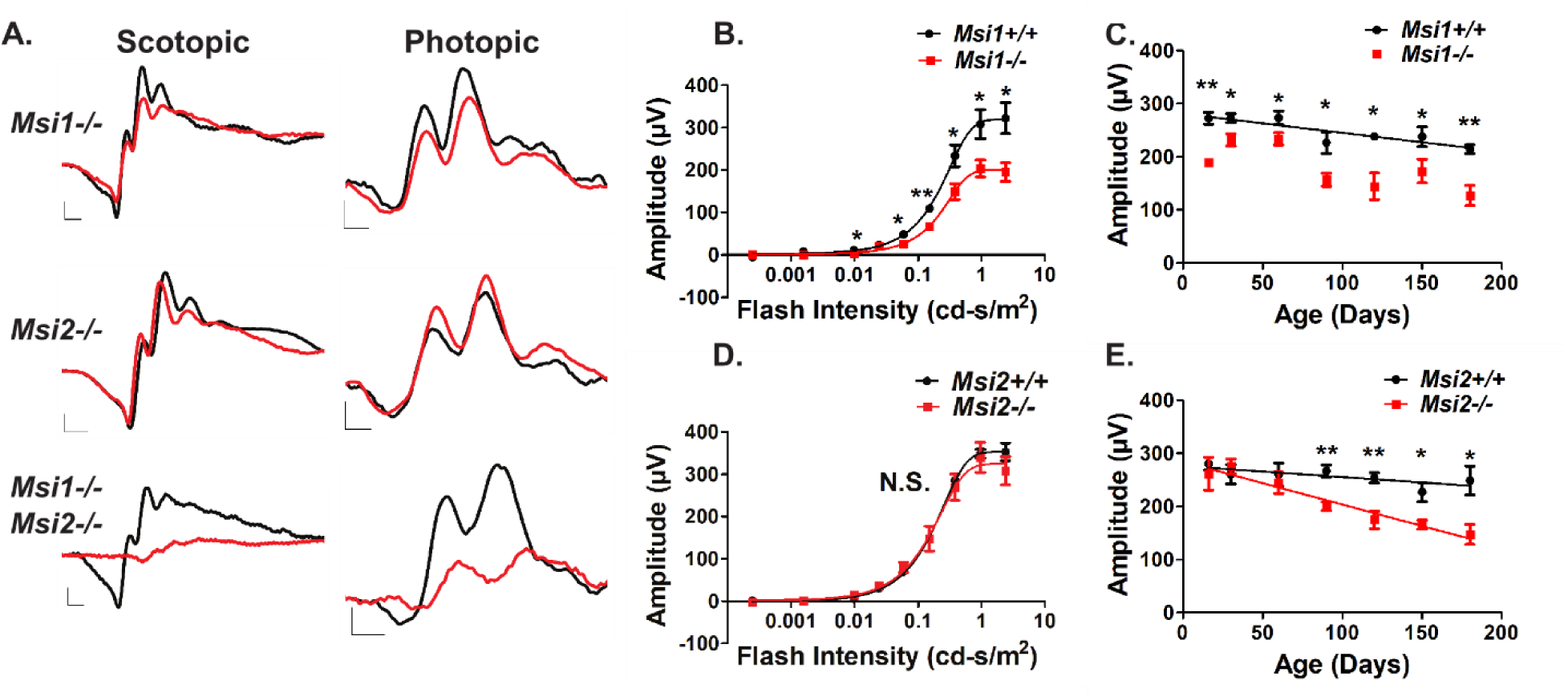
Rod cell specific defect of the double *Msi1* and *Msi2* knockout. **A.** Representative scotopic and photopic electroretinograms (ERGs) from the *rod-Msi1-/-*, *rod-Msi2-/-*, and *rod-Msi1-/-: Msi2-/-* mice at PN16. Scotopic ERGs were obtained after overnight dark adaptation using 0.151 cd-s/m2 flashes while photopic ERGs were obtained with 7.6 cd-s/m2 flashes under light-adapted conditions using a rod-saturating white background light (Scotopic scale bar: x-axis = 10ms, y-axis = 100 µV; Photopic scale bar: x-axis = 10ms, y-axis = 20 µV). **B.** Intensity response plot of the scotopic “a”-wave from *rod-Msi1-/-* mice (*=P-value < 0.05; **=P-value < 0.01; ***=P-value < 0.001). **C.** Plot of the rod photoreceptor “a”-wave response from *rod-Msi1-/-* mice against the age of the mouse during which the ERG was recorded. **D.** Intensity response plot of the scotopic “a”-wave response from *rod-Msi2-/-* mice (*=P-value < 0.05; **=P-value < 0.01; ***=P-value < 0.001). **E.** Plot of the rod photoreceptor “a”-wave response from *rod-Msi2-/-* mice against the age of the mouse during which the ERG was recorded. All data is shown as the mean ± the SEM, and statistical analyses were carried out using the homoscedastic unpaired student’s t-test (*=P<0.05).

### Progressive neuronal degeneration in the absence of the Musashi proteins

We next wanted to examine the mechanism behind the photoreceptor dysfunction seen in the *ret-Msi1-/-:Msi2-/-* mouse model. One of the common causes of a reduced ERG is photoreceptor cell death. Therefore, we performed histological analysis of the *ret-Msi1-/-:Msi2-/-* mice at PN5, PN10, PN16, and PN180 (Figure 5A–D). In *ret-Msi1-/-:Msi2-/-* mice at PN5, even before the neural retina has completely differentiated, there is a reduction in the neuroblast layer (NBL) thickness which was quantified across the superior-inferior axis (Figure 5A, left and right panels). There is also a more disordered arrangement of NBL nuclei in *ret-Msi1-/-:Msi2-/-* mice with cells more tightly packed together compared to its littermate control (Figure 5A, left panel). At PN10, the outer nuclear layer (ONL), inner nuclear layer (INL), and ganglion cell layer (GCL) of the retina all form in *ret-Msi1-/-:Msi2-/-* mice but there is a reduction in the number of layers of photoreceptor nuclei (Figure 5B, left and right panels). At PN16, the number of layers of ONL nuclei continue to decrease suggesting that photoreceptor cell death is occurring (Figure 5C, left and middle panels). However, at this age, there are no statistically significant changes in the number of layers of INL nuclei (Figure 5C, left and right panels). By 6 months of age, the retina of *ret-Msi1-/-:Msi2-/-* mice was severely degenerated with a complete loss of ONL nuclei in addition to a significant reduction in the number of layers of INL nuclei (Figure 5D, left, middle, and right panels).

**Figure 5:**
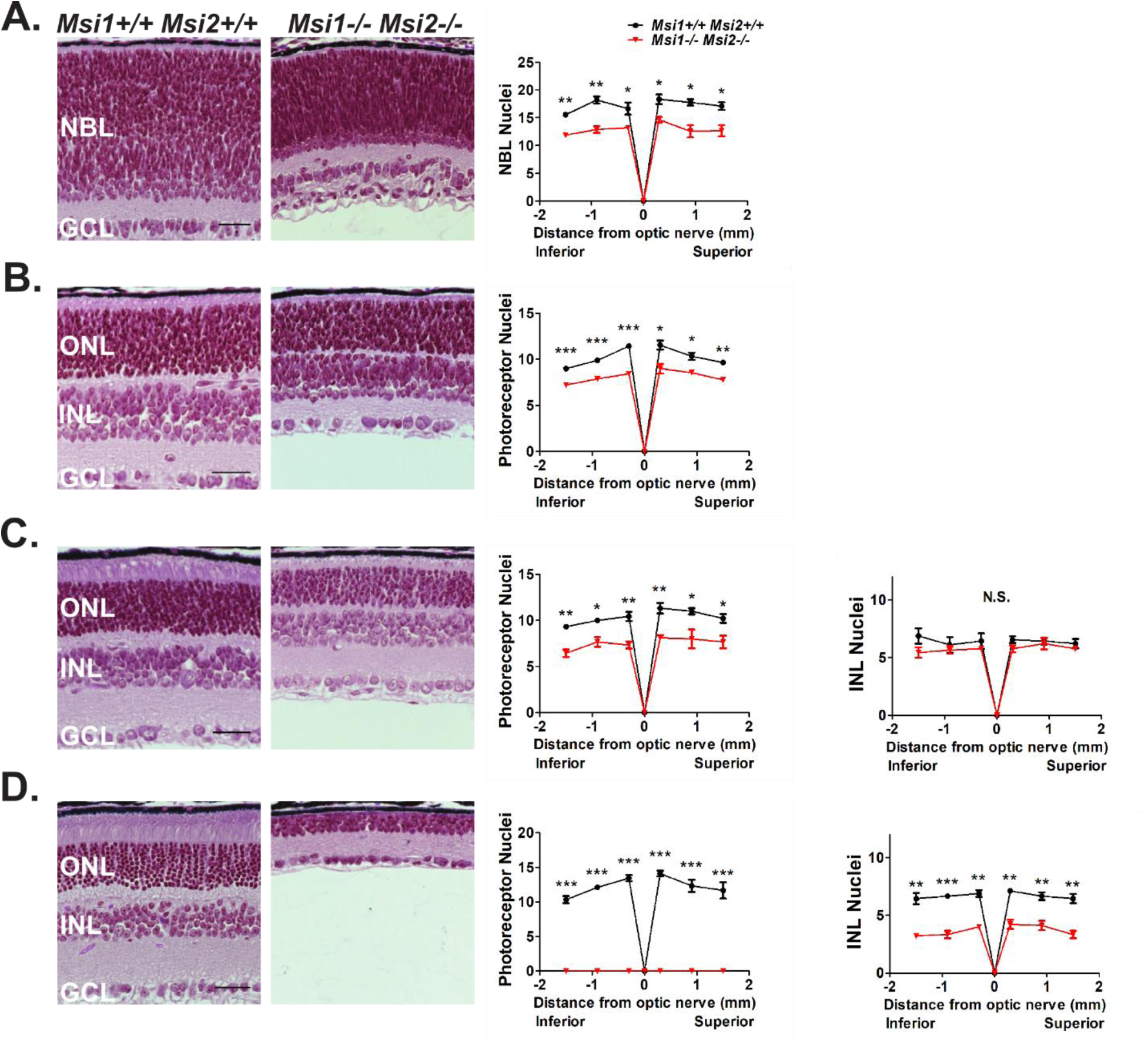
Retinal cell death occurs in the absence of the Musashi proteins. Left: Brightfield microscopic images of H&E stained retinal cross sections from the *ret-Msi1-/-: Msi2-/-* mice at PN5 **(A),** PN10 **(B)**, PN16 **(C)**, and PN180 **(D)**. Right: Spider plot of the indicated layer thickness at six regions from the inferior to superior retina in the *ret-Msi1-/-: Msi2-/-* mice at PN5 **(A),** PN10 **(B)**, PN16 **(C)**, and PN180 **(D)** (NBL: neuroblast layer, ONL: outer nuclear layer, INL: inner nuclear layer, and GCL: ganglion cell layer). All data is shown as the mean ± the SEM, and statistical analyses were carried out using the homoscedastic unpaired student’s t-test (*=P-value < 0.05; **=P-value < 0.01; ***=P-value < 0.001).

### The Musashi proteins are crucial for photoreceptor outer segment and axoneme development

Photoreceptor cells are present in the *ret-Msi1-/-:Msi2-/-* as indicated by the well-defined ONL (Figure 1C). We therefore examined the structure of the OS in *ret-Msi1-/-:Msi2-/-* mice at PN16 by immunofluorescence microscopy using three different OS markers, anti-Peripherin-2 (PRPH2: OS marker), anti-Phosphodiesterase-6β (PDE6β: rod OS marker), and peanut agglutinin (PNA: cone OS marker). After staining retinal cross sections from *ret-Msi1-/-:Msi2-/-* mice with PRPH2 and PNA, we observed a severe shortening of the photoreceptor outer segment (Figure 6A). This result was not limited to PRPH2, as staining with the rod OS marker PDE6β demonstrated the same phenotype (Figure 6B). The outer segment of cone photoreceptors also appears to be severely shortened as shown by the abnormal PNA staining (Figure 6A–B). Lastly, no mislocalization of PDE6β or PRPH2 is found in the ONL or inner segment of *ret-Msi1-/-:Msi2-/-* mice suggesting that while the Musashi proteins are required for outer segment formation they are not regulating trafficking or localization of OS-resident proteins (Figure 6B).

**Figure 6:**
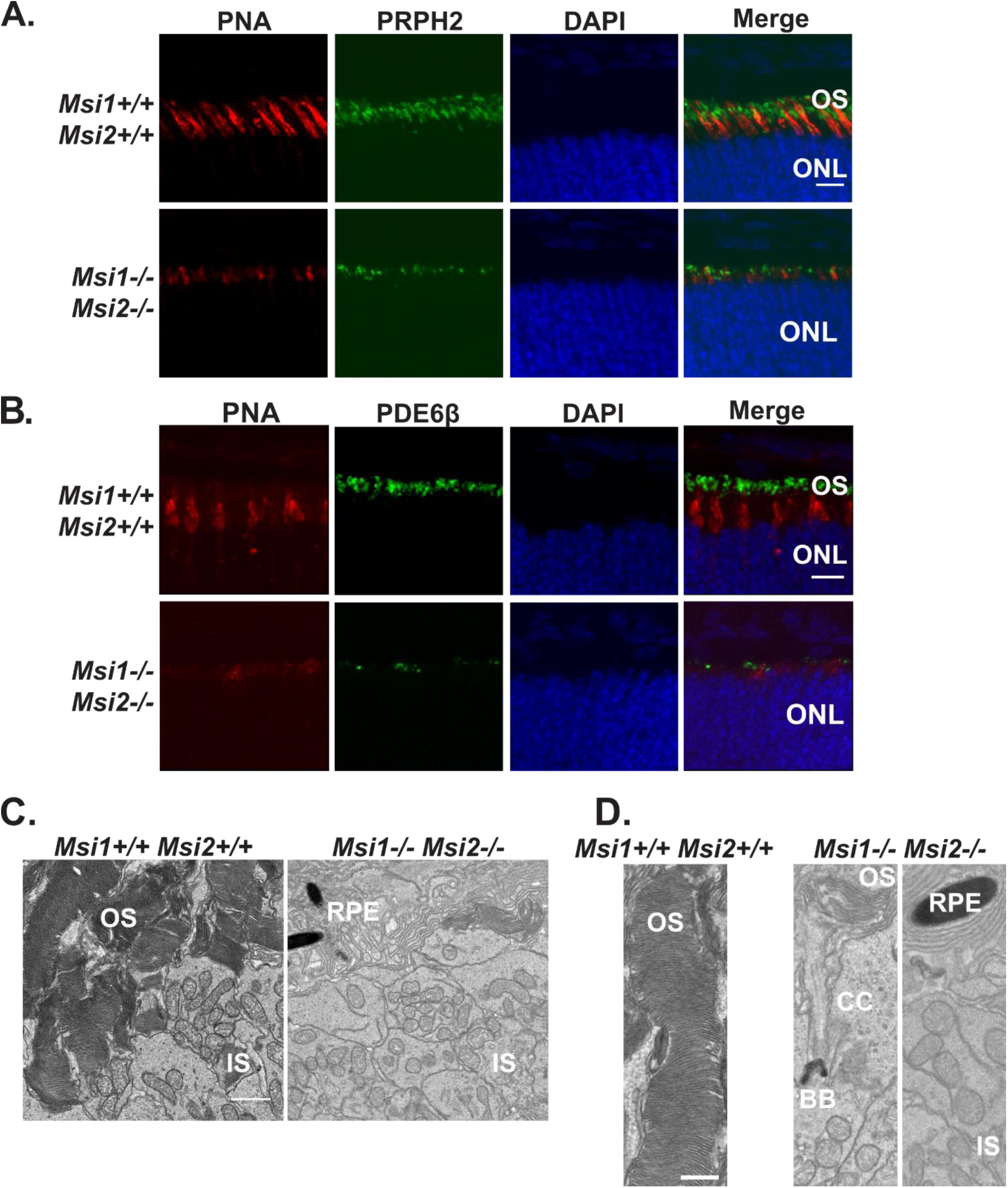
Abnormal development of OS in the absence of MSI1 and MSI2. **A.** Immunofluorescence microscopy images of retinal cross sections from the *ret-Msi1-/-: Msi2-/-* mice at PN10 stained with anti-peripherin-2 antibody (PRPH2: OS marker - Green) and peanut agglutinin (PNA: cone OS marker - Red) along with a DAPI nuclear counterstain (Blue). Scale bar = 20 µm. **B.** Immunofluorescence microscopy images of retinal cross sections from the *ret-Msi1-/-: Msi2-/-* mice at PN10 stained with anti-phosphodiesterase-6β antibody (PDE6β: rod OS marker - Green) and peanut agglutinin (PNA: cone OS marker - Red) along with a DAPI counterstain (Blue). (OS: outer segment and ONL: outer nuclear layer). Scale bar = 20 µm. **C.** Low magnification transmission electron microscopy images of ultrathin retinal sections from *ret-Msi1-/-: Msi2-/-* mice at PN10 visualizing the boundary between the OS and IS showing the lack of typical outer segments in the absence of the Musashi proteins (OS: outer segment, IS: inner segment, and RPE: retinal pigment epithelium). Scale bar = 2 µm. **D.** High magnification transmission electron microscopy images of ultrathin retinal sections from *ret-Msi1-/-: Msi2-/-* mice at PN10 visualizing the boundary between the OS and IS showing that the OS either does not form (far right) or is dysmorphic (middle) in the absence of the Musashi proteins (OS: outer segment, CC: connecting cilium, BB: basal body, RPE: retinal pigment epithelium, and IS: inner segment). Scale bar = 1 µm.

Using transmission electron microscopy, we imaged ultrathin retinal sections from *ret-Msi1-/-:Msi2-/-* mice at PN10 when the OS begins to elaborate. When examining the OS/IS boundary in *ret-Msi1-/-:Msi2-/-* mice by electron microscopy, we observed very little, if any, conspicuous OS (Figure 6C). Instead, the IS of the *ret-Msi1-/-:Msi2-/-* mice appears to come in direct contact with the RPE (Figure 6C–D). At higher magnification, the photoreceptors of *ret-Msi1-/-:Msi2-/-* mice displayed either no OS or aberrant and undersized OS (Figure 6D left, middle, and right panels).

To examine the structure of the connecting cilium and the axoneme, we stained retinal cross sections from *ret-Msi1-/-:Msi2-/-* mice at PN10 using antibodies directed against the established markers of murine connecting cilium (glutamylated and acetylated tubulin) and axoneme (MAK) (29–32). Probing with glutamylated and acetylated α-tubulin antibodies showed that there were no changes in the length of the CC (Figure 7A, C-D). Contrarily, staining with the anti-MAK antibody showed a substantial reduction in the length of the axoneme accompanied with punctate staining suggesting a severe structural defect of the axoneme (Figure 7A–B).

**Figure 7:**
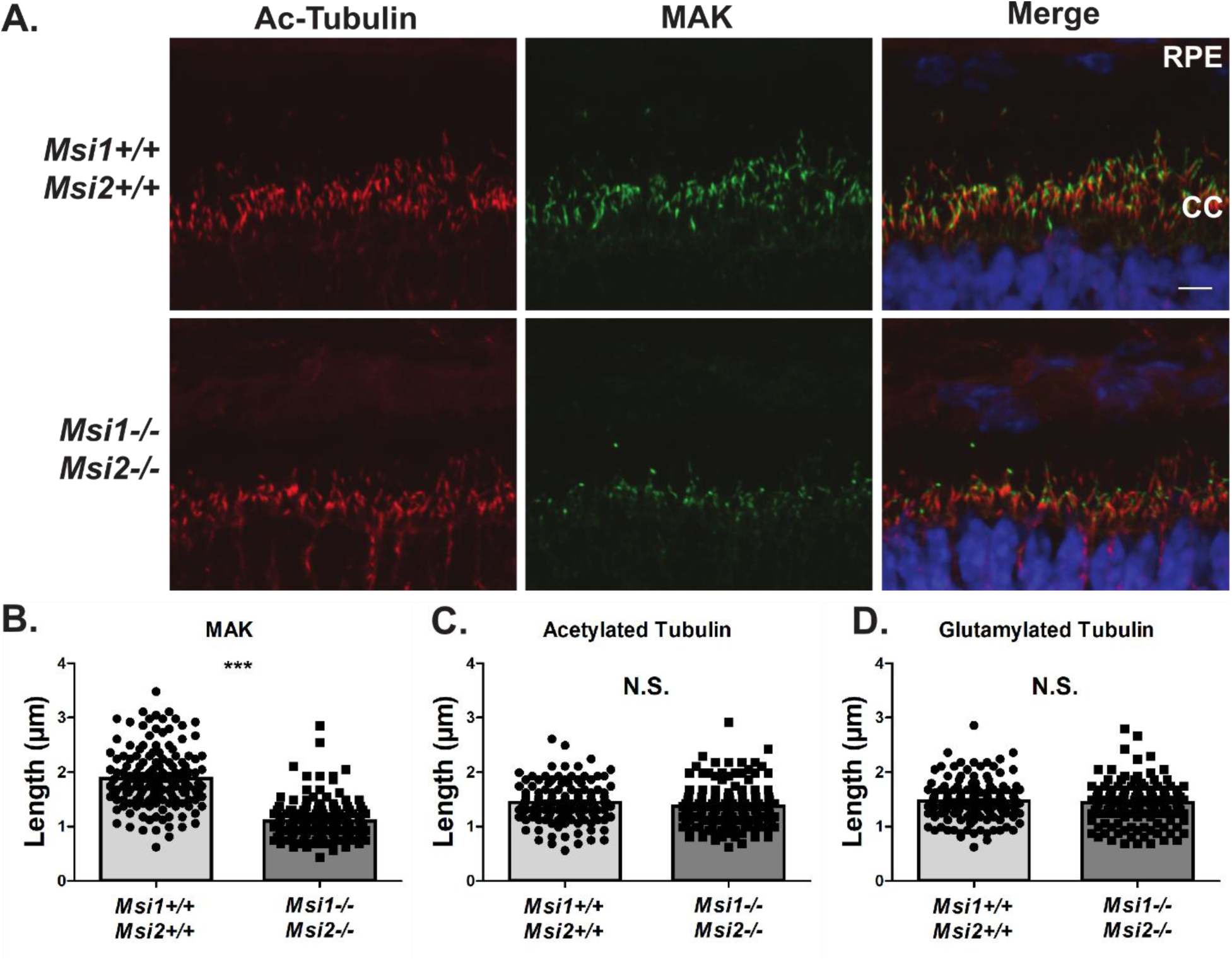
The Musashi proteins are crucial for photoreceptor axoneme development. **A.** Immunofluorescence microscopy images of retinal cross sections from the *ret-Msi1-/-: Msi2-/-* mice at PN10 stained with acetylated-α-tubulin antibody (Ac-Tubulin: Red) and male germ cell-associated kinase antibody (MAK: Green) along with DAPI counterstain (Blue) (RPE: retinal pigment epithelium, CC: connecting cilium, and ONL: outer nuclear layer). Scatter bar plot showing the distribution of length measurements for the photoreceptor axoneme by MAK staining **(B)** and connecting cilium by Ac-tubulin staining **(C)** and glutamylated tubulin staining **(D)**. Retinal sections were obtained from PN10 musashi knockouts and littermate controls.

### The Musashi proteins promote splicing of photoreceptor specific exons

Our previous studies suggested that the Musashi proteins are regulating alternative splicing of their target pre-mRNAs in vertebrate photoreceptors (8). To test if the Musashi proteins are responsible for the inclusion of photoreceptor specific exon, we analyzed the splicing in *ret-Msi1-/-:Msi2-/-* mice of pre-mRNAs from cilia-and OS-related genes that we previously showed to express photoreceptor specific isoforms (Figure 8). We witnessed a drastic reduction in alternative exon inclusion in *ret-Msi1-/-:Msi2-/-* mice for all tested transcripts (Figure 8A). We also analyzed isoform expression at the protein level for TTC8 (Tetratricopeptide repeat domain 8) since we had an antibody that specifically recognizes the photoreceptor-specific isoform. TTC8 also referred as Bardet-Biedl Syndrome Protein (BBS8) is part of the BBSome complex that is known play an important role in photoreceptor outer segment morphogenesis (33, 34). We used two different antibodies, a pan-antibody that recognizes all TTC8 protein isoforms (Pan-TTC8) and the other that recognizes the photoreceptor-specific isoform of Ttc8 by binding the epitope encoded by Exon 2A (the photoreceptor-specific exon of *Ttc8*) (Figure 8B). After probing retinal lysates from the *ret-Msi1-/-:Msi2-/-* mice with the pan-TTC8 antibody, we observed faster migration of the TTC8 protein compared to the littermate control suggesting that the Exon 2A was not included (Figure 8B). Concordantly, when probing for the photoreceptor-specific isoform of TTC8 using the Ttc8 Exon 2A antibody, we saw the absence of this isoform in *ret-Msi1-/-:Msi2-/-* mice (Figure 8B). Taken together, these results demonstrate that the Musashi proteins are required for the inclusion of photoreceptor specific alternative exons.

**Figure 8:**
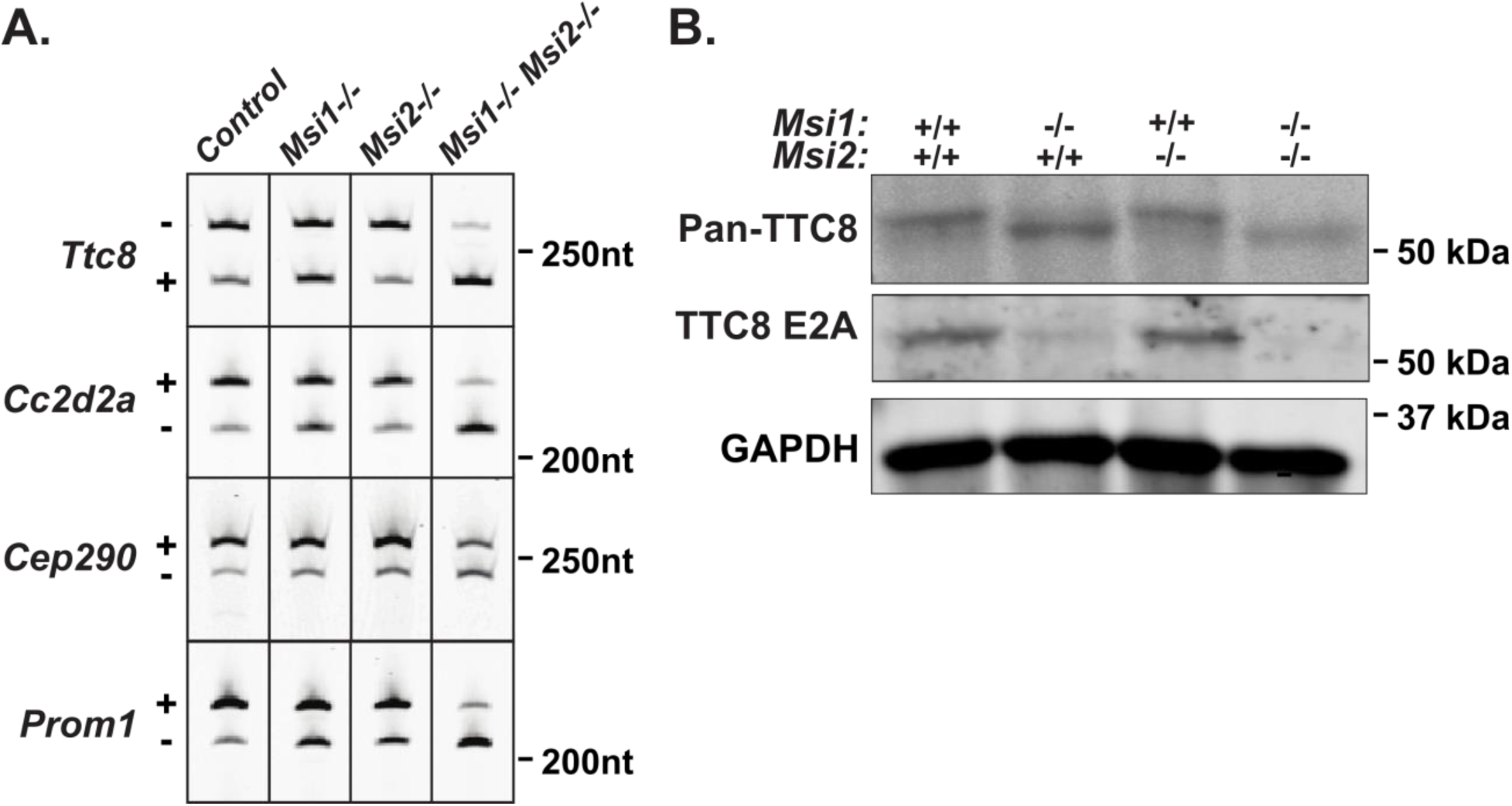
The Musashi proteins regulate alternative splicing of their target transcripts. **A.** Reverse transcriptase PCR splicing assay using total RNA purified from retinal lysates of *ret-Msi1-/-*, *ret-Msi2-/-*, and *ret-Msi1-/-: Msi2-/-* mice. *Ttc8*, *Cc2d2a*, *Cep290*, and *Prom1* are four cilia- and OS- related transcripts shown to have reduced photoreceptor-specific exon inclusion in the absence of MSI1 and MSI2. **B.** Immunoblot of retinal lysates from *ret-Msi1-/-*, *ret-Msi2-/-*, and *ret-Msi1-/-: Msi2-/-* mice. After probing with the pan-TTC8 antibody (top), a change in the migration of the TTC8 protein is observed in the absence of MSI1 and MSI2 suggesting that the peptide encoded by Exon 2A was not included. When probing with the TTC8 E2A antibody (middle), photoreceptor-specific isoform of TTC8 was not observed in the absence of MSI1 and MSI2.

## DISCUSSION

### MSI1 and MSI2 are required for photoreceptor morphogenesis but not specification

Our data shows the requirement for MSI1 and MSI2 in photoreceptor cells. Double knockout of *Msi1* and *Msi2* in retinal progenitors results in complete loss of vision. Two lines of evidence demonstrate that this loss of vision is due to a defect in photoreceptor morphogenesis, rather than early developmental defects. First, the specification of retinal progenitors to photoreceptor cells was not affected by loss of Musashi. The retina of the knockout mice had laminated nuclear layers indicating normal development of the retina. The photoreceptor cells retained their characteristic morphology and expressed cell type specific proteins such as peripherin and PDE6. Importantly, removal of *Msi1* and *Msi2* in rod photoreceptors driven by *Nrl-Cre* caused loss of scotopic photoresponse. Thus, the vision phenotype is not due to impairment of the early stages of retinal development and is caused by a defect specific to photoreceptor cells.

Morphological examination by electron microscopy and immunofluorescence showed that the outer segment of the photoreceptors lacking Musashi is either missing or is stunted and disorganized. In addition, axoneme was shortened. In contrast, the connecting cilium has normal length and did not have obvious defects. Trafficking of PDE6 and peripherin through the connecting cilium also appears to be unaffected and the two proteins localize to the stunted outer segment wherever one is present. Taken together our findings demonstrates a requirement for Musashi in the morphogenesis and function of the photoreceptor outer segment that appears not to affect protein trafficking.

### Musashi is needed for inclusion of photoreceptor-specific exons

RT-PCR analysis of alternative splicing in the retina of *Msi1* and *Msi2* knockout mice showed that inclusion of photoreceptor specific exons in the mature transcripts is dependent on the Musashi proteins. We confirmed this finding using immunoblotting with antibody that recognizes photoreceptor-specific isoform of TTC8. The effect of *Msi1* knockout on splicing is stronger compared to Msi2 knockdown. It remains to be determined if this observation reflects a dominant role for MSI1 in splicing control, or derives from to the timing of the embryonic knockout of the two genes relative to the postnatal developmental switch from Msi1 to Msi2 expression in the retina. Our data demonstrates for the first time that Musashi regulates splicing *in vivo* and impacts dramatically the inclusion levels of the exons it controls. This is a novel role for Musashi that is distinct from it known function in controlling translation in the cytosol.

### Functional redundancy and developmental switch within the Musashi protein family

In vertebrates, the Musashi protein family consists of two paralogues, MSI1 and MSI2, which have high degree of sequence identity, and have arisen from a gene duplication event (35, 36). The RNA binding domains of MSI1 and MSI2 have approximately 90% sequence identity and recognize the same UAG sequence motif *in vitro* and *in vivo* (37–40). The high degree of similarity suggest that the two proteins are likely to be functionally redundant when co-expressed in the same cells. Indeed, we observed only minor reductions in visual function after the loss of either MSI1 or MSI2 alone whereas the combined loss of MSI1 and MSI2 resulted in a complete loss of visual function (Figure 2). Similarly, inclusion of photoreceptor specific exons is promoted by both proteins, and the double knockout produces stronger effect on splicing than the knockouts of either *Msi1* or *Msi2*. The functional redundancy in photoreceptor cells that we observe is in agreement with previous reports of redundancy between MSI1 and MSI2 in other cell types (12, 13).

Despite the proposed functional redundancy between the two Musashi proteins the phenotype of the single *Msi1* and *Msi2* knockouts show distinct progression of vision loss. *Msi1* knockouts have reduced vision at birth, followed by minor decline as the animals age. This decline is unlikely to be associated with the lack of Musashi, as it tracks the normal reduction in visual response observed in the wild type controls. In contrast, *Msi2* knockouts do not show significant visual defect at the time of eye opening (P16), but their vision progressively deteriorates with age. This difference in the phenotypes can be explained by the developmental timing of the MSI1 and MSI2 protein expression. A burst in MSI1 protein expression precedes the critical period for rod photoreceptor outer segment morphogenesis between birth and eye opening and MSI1 levels remain high until the eyes open at P16. The MSI1 expression begins a gradual decline at P13 and the MSI1 protein is replaced by increase in MSI2 levels. This data shows distinct roles for MSI1 and MSI2 in photoreceptor morphogenesis and photoreceptor maintenance, respectively. The developmental switch we observe raises the question of potential functional differences in the two Musashi protein, that require MSI1 expression during photoreceptor morphogenesis and MSI2 for photoreceptor maintenance.

Our work highlights roles for MSI1 and MSI2 in photoreceptor morphogenesis and survival. An interesting aspect of the function of the Musashi proteins in the retina is their apparently mutually exclusive roles at different stages of development. At early stages of development, MSI1 and MSI2 support the renewal and proliferation of retinal precursor cells. At late stages of retinal development and in the adult retina MSI1 and MSI2 are required for morphogenesis of the differentiated photoreceptor cells and survival of mature neurons. Our studies point to the need for MSI in controlling the alternative splicing in photoreceptor cells. It is important to note that the canonical function of the Musashi proteins is to control mRNA translation in the cytosol (41, 42), where they can either block or enhance translation of mRNA depending on cellular context (43–48). Future studies will be aimed at determining the mechanism(s) for the need for Musashi in vision and the regulation of the developmental switch between MSI1 and MSI2.

## Acknowledgments

The authors thank Maxim Sokolov, John Hollander, Ronald Gross for their feedback on the work. We also thank Dr. Christopher Lengner for the generous donation of the *Msi1fl/fl Msi2fl/fl* mice and Dr. Andrew Goldberg for PRPH2 antibody.

## AUTHOR CONTRIBUTIONS

P.S. and V.R jointly conceived and supervised this study and edited the manuscript. J.S. designed and performed experiments and wrote the manuscript. F.M and B.J. designed and performed experiments.

## REFERENCES

1. Grau-Bové, X., Ruiz-Trillo, I., and Irimia, M. (2018) Origin of exon skipping-rich transcriptomes in animals driven by evolution of gene architecture. Genome Biol. 19, 135

2. Li, Y. I., Sanchez-Pulido, L., Haerty, W., and Ponting, C. P. (2014) RBFOX and PTBP1 proteins regulate the alternative splicing of micro-exons in human brain transcripts. Genome Res. 10.1101/gr.181990.114

3. Irimia, M., Weatheritt, R. J., Ellis, J. D., Parikshak, N. N., Gonatopoulos-Pournatzis, T., Babor, M., Quesnel-Vallières, M., Tapial, J., Raj, B., O’Hanlon, D., Barrios-Rodiles, M., Sternberg, M. J. E., Cordes, S. P., Roth, F. P., Wrana, J. L., Geschwind, D. H., and Blencowe, B. J. (2014) A Highly Conserved Program of Neuronal Microexons Is Misregulated in Autistic Brains. Cell. 159, 1511–1523

4. Jensen, K. B., Dredge, B. K., Stefani, G., Zhong, R., Buckanovich, R. J., Okano, H. J., Yang, Y. Y. L., and Darnell, R. B. (2000) Nova-1 Regulates Neuron-Specific Alternative Splicing and Is Essential for Neuronal Viability. Neuron. 25, 359–371

5. Gehman, L. T., Stoilov, P., Maguire, J., Damianov, A., Lin, C.-H., Shiue, L., Ares, M., Mody, I., and Black, D. L. (2011) The splicing regulator Rbfox1 (A2BP1) controls neuronal excitation in the mammalian brain. Nat Genet. 43, 706–711

6. Ule, J., Ule, A., Spencer, J., Williams, A., Hu, J.-S., Cline, M., Wang, H., Clark, T., Fraser, C., Ruggiu, M., Zeeberg, B. R., Kane, D., Weinstein, J. N., Blume, J., and Darnell, R. B. (2005) Nova regulates brain-specific splicing to shape the synapse. Nat. Genet. 37, 844–852

7. Vuong, C. K., Wei, W., Lee, J.-A., Lin, C.-H., Damianov, A., de la Torre-Ubieta, L., Halabi, R., Otis, K. O., Martin, K. C., O’Dell, T. J., and Black, D. L. (2018) Rbfox1 Regulates Synaptic Transmission Through the Inhibitory Neuron Specific vSNARE Vamp1. Neuron. 98, 127–141.e7

8. Murphy, D., Cieply, B., Carstens, R., Ramamurthy, V., and Stoilov, P. (2016) The Musashi 1 Controls the Splicing of Photoreceptor-Specific Exons in the Vertebrate Retina. PLOS Genet. 12, e1006256

9. Ling, J. P., Wilks, C., Charles, R., Leavey, P. J., Ghosh, D., Jiang, L., Santiago, C. P., Pang, B., Venkataraman, A., Clark, B. S., Nellore, A., Langmead, B., and Blackshaw, S. (2020) ASCOT identifies key regulators of neuronal subtype-specific splicing. Nat. Commun. 11, 1–12

10. Ohyama, T., Nagata, T., Tsuda, K., Kobayashi, N., Imai, T., Okano, H., Yamazaki, T., and Katahira, M. (2012) Structure of Musashi1 in a complex with target RNA: the role of aromatic stacking interactions. Nucleic Acids Res. 40, 3218–3231

11. Sakakibara, S., Nakamura, Y., Satoh, H., and Okano, H. (2001) RNA-Binding Protein Musashi 2: Developmentally Regulated Expression in Neural Precursor Cells and Subpopulations of Neurons in Mammalian CNS. J. Neurosci. 21, 8091–8107

12. Sakakibara, S., Nakamura, Y., Yoshida, T., Shibata, S., Koike, M., Takano, H., Ueda, S., Uchiyama, Y., Noda, T., and Okano, H. (2002) RNA-binding protein Musashi family: Roles for CNS stem cells and a subpopulation of ependymal cells revealed by targeted disruption and antisense ablation. Proc. Natl. Acad. Sci. 99, 15194–15199

13. Li, N., Yousefi, M., Nakauka-Ddamba, A., Li, F., Vandivier, L., Parada, K., Woo, D.-H., Wang, S., Naqvi, A. S., Rao, S., Tobias, J., Cedeno, R. J., Minuesa, G., Y, K., Barlowe, T. S., Valvezan, A., Shankar, S., Deering, R. P., Klein, P. S., Jensen, S. T., Kharas, M. G., Gregory, B. D., Yu, Z., and Lengner, C. J. (2015) The Msi Family of RNA-Binding Proteins Function Redundantly as Intestinal Oncoproteins. Cell Rep. 13, 2440–2455

14. Pearring, J. N., Salinas, R. Y., Baker, S. A., and Arshavsky, V. Y. (2013) Protein sorting, targeting and trafficking in photoreceptor cells. Prog. Retin. Eye Res. 36, 24–51

15. Riazuddin, S. A., Iqbal, M., Wang, Y., Masuda, T., Chen, Y., Bowne, S., Sullivan, L. S., Waseem, N. H., Bhattacharya, S., Daiger, S. P., Zhang, K., Khan, S. N., Riazuddin, S., Hejtmancik, J. F., Sieving, P. A., Zack, D. J., and Katsanis, N. (2010) A Splice-Site Mutation in a Retina-Specific Exon of BBS8 Causes Nonsyndromic Retinitis Pigmentosa. Am. J. Hum. Genet. 86, 805–812

16. Murphy, D., Singh, R., Kolandaivelu, S., Ramamurthy, V., and Stoilov, P. (2015) Alternative Splicing Shapes the Phenotype of a Mutation in BBS8 To Cause Nonsyndromic Retinitis Pigmentosa. Mol. Cell. Biol. 35, 1860–1870

17. Rachel, R. A., Li, T., and Swaroop, A. (2012) Photoreceptor sensory cilia and ciliopathies: focus on CEP290, RPGR and their interacting proteins. Cilia. 1, 22

18. Veleri, S., Manjunath, S. H., Fariss, R. N., May-Simera, H., Brooks, M., Foskett, T. A., Gao, C., Longo, T. A., Liu, P., Nagashima, K., Rachel, R. A., Li, T., Dong, L., and Swaroop, A. (2014) Ciliopathy-associated gene Cc2d2a promotes assembly of subdistal appendages on the mother centriole during cilia biogenesis. Nat. Commun. 5, 1–12

19. Zacchigna, S., Oh, H., Wilsch-Bräuninger, M., Missol-Kolka, E., Jászai, J., Jansen, S., Tanimoto, N., Tonagel, F., Seeliger, M., Huttner, W. B., Corbeil, D., Dewerchin, M., Vinckier, S., Moons, L., and Carmeliet, P. (2009) Loss of the Cholesterol-Binding Protein Prominin-1/CD133 Causes Disk Dysmorphogenesis and Photoreceptor Degeneration. J. Neurosci. 29, 2297–2308

20. Ba-Abbad, R., Arno, G., Carss, K., Stirrups, K., Penkett, C. J., Moore, A. T., Michaelides, M., Raymond, F. L., Webster, A. R., and Holder, G. E. (2016) Mutations in CACNA2D4 Cause Distinctive Retinal Dysfunction in Humans. Ophthalmology. 123, 668–671.e2

21. Johnson, J., Fremeau, R. T., Duncan, J. L., Rentería, R. C., Yang, H., Hua, Z., Liu, X., LaVail, M. M., Edwards, R. H., and Copenhagen, D. R. (2007) Vesicular Glutamate Transporter 1 Is Required for Photoreceptor Synaptic Signaling But Not For Intrinsic Visual Functions. J. Neurosci. 27, 7245–7255

22. A simple polymerase chain reaction assay for genotyping the retinal degeneration mutation (Pdebrd1) in FVB/N-derived transgenic mice (2001) Lab. Anim. 35, 153–156

23. Pak, J. S., Lee, E.-J., and Craft, C. M. (2015) The retinal phenotype of Grk1−/− is compromised by a Crb1rd8 mutation. Mol. Vis. 21, 1281–1294

24. Wright, Z. C., Singh, R. K., Alpino, R., Goldberg, A. F. X., Sokolov, M., and Ramamurthy, V. (2016) ARL3 regulates trafficking of prenylated phototransduction proteins to the rod outer segment. Hum. Mol. Genet. 25, 2031–2044

25. Wright, Z. C., Loskutov, Y., Murphy, D., Stoilov, P., Pugacheva, E., Goldberg, A. F. X., and Ramamurthy, V. (2018) ADP-Ribosylation Factor-Like 2 (ARL2) regulates cilia stability and development of outer segments in rod photoreceptor neurons. Sci. Rep. 8, 1–12

26. Furuta, Y., Lagutin, O., Hogan, B. L. M., and Oliver, G. C. (2000) Retina- and ventral forebrain-specific Cre recombinase activity in transgenic mice. genesis. 26, 130–132

27. Guan, W., Cao, J.-W., Liu, L.-Y., Zhao, Z.-H., Fu, Y., and Yu, Y.-C. (2017) Eye opening differentially modulates inhibitory synaptic transmission in the developing visual cortex. eLife. 6, e32337

28. Brightman, D. S., Razafsky, D., Potter, C., Hodzic, D., and Chen, S. (2016) Nrl-Cre transgenic mouse mediates loxP recombination in developing rod photoreceptors. Genes. N. Y. N 2000. 54, 129–135

29. Dilan, T. L., Moye, A. R., Salido, E. M., Saravanan, T., Kolandaivelu, S., Goldberg, A. F. X., and Ramamurthy, V. (2019) ARL13B, a Joubert Syndrome-Associated Protein, Is Critical for Retinogenesis and Elaboration of Mouse Photoreceptor Outer Segments. J. Neurosci. 39, 1347–1364

30. Arikawa, K., and Williams, D. S. (1993) Acetylated alpha-tubulin in the connecting cilium of developing rat photoreceptors. Invest. Ophthalmol. Vis. Sci. 34, 2145–2149

31. Grau, M. B., Masson, C., Gadadhar, S., Rocha, C., Tort, O., Sousa, P. M., Vacher, S., Bieche, I., and Janke, C. (2017) Alterations in the balance of tubulin glycylation and glutamylation in photoreceptors leads to retinal degeneration. J. Cell Sci. 130, 938–949

32. Omori, Y., Chaya, T., Katoh, K., Kajimura, N., Sato, S., Muraoka, K., Ueno, S., Koyasu, T., Kondo, M., and Furukawa, T. (2010) Negative regulation of ciliary length by ciliary male germ cell-associated kinase (Mak) is required for retinal photoreceptor survival. Proc. Natl. Acad. Sci. 107, 22671–22676

33. Hsu, Y., Garrison, J. E., Kim, G., Schmitz, A. R., Searby, C. C., Zhang, Q., Datta, P., Nishimura, D. Y., Seo, S., and Sheffield, V. C. (2017) BBSome function is required for both the morphogenesis and maintenance of the photoreceptor outer segment. PLoS Genet. 10.1371/journal.pgen.1007057

34. Dilan, T. L., Singh, R. K., Saravanan, T., Moye, A., Goldberg, A. F. X., Stoilov, P., and Ramamurthy, V. (2018) Bardet–Biedl syndrome-8 (BBS8) protein is crucial for the development of outer segments in photoreceptor neurons. Hum. Mol. Genet. 27, 283–294

35. Ohyama, T., Nagata, T., Tsuda, K., Kobayashi, N., Imai, T., Okano, H., Yamazaki, T., and Katahira, M. (2012) Structure of Musashi1 in a complex with target RNA: the role of aromatic stacking interactions. Nucleic Acids Res. 40, 3218–3231

36. Sutherland, J. M., Siddall, N. A., Hime, G. R., and McLaughlin, E. A. (2015) RNA binding proteins in spermatogenesis: an in depth focus on the Musashi family. Asian J. Androl. 17, 529–536

37. Uren, P. J., Vo, D. T., Araujo, P. R. de, Pötschke, R., Burns, S. C., Bahrami-Samani, E., Qiao, M., Abreu, R. de S., Nakaya, H. I., Correa, B. R., Kühnöl, C., Ule, J., Martindale, J. L., Abdelmohsen, K., Gorospe, M., Smith, A. D., and Penalva, L. O. F. (2015) RNA-Binding Protein Musashi1 Is a Central Regulator of Adhesion Pathways in Glioblastoma. Mol. Cell. Biol. 35, 2965–2978

38. Bennett, C. G., Riemondy, K., Chapnick, D. A., Bunker, E., Liu, X., Kuersten, S., and Yi, R. (2016) Genome-wide analysis of Musashi-2 targets reveals novel functions in governing epithelial cell migration. Nucleic Acids Res. 44, 3788–3800

39. Rentas, S., Holzapfel, N. T., Belew, M. S., Pratt, G. A., Voisin, V., Wilhelm, B. T., Bader, G. D., Yeo, G. W., and Hope, K. J. (2016) Musashi-2 attenuates AHR signalling to expand human haematopoietic stem cells. Nature. 532, 508–511

40. Lan, L., Xing, M., Douglas, J. T., Gao, P., Hanzlik, R. P., and Xu, L. (2017) Human oncoprotein Musashi-2 N-terminal RNA recognition motif backbone assignment and identification of RNA-binding pocket. Oncotarget. 8, 106587–106597

41. Kudinov, A. E., Karanicolas, J., Golemis, E. A., and Boumber, Y. (2017) Musashi RNA-Binding Proteins as Cancer Drivers and Novel Therapeutic Targets. Clin. Cancer Res. 23, 2143–2153

42. Fox, R. G., Park, F. D., Koechlein, C. S., Kritzik, M., and Reya, T. (2015) Musashi Signaling in Stem Cells and Cancer. Annu. Rev. Cell Dev. Biol. 31, 249–267

43. Imai, T., Tokunaga, A., Yoshida, T., Hashimoto, M., Mikoshiba, K., Weinmaster, G., Nakafuku, M., and Okano, H. (2001) The Neural RNA-Binding Protein Musashi1 Translationally Regulates Mammalian numb Gene Expression by Interacting with Its mRNA. Mol. Cell. Biol. 21, 3888–3900

44. Battelli, C., Nikopoulos, G. N., Mitchell, J. G., and Verdi, J. M. (2006) The RNA-binding protein Musashi-1 regulates neural development through the translational repression of p21WAF-1. Mol. Cell. Neurosci. 31, 85–96

45. Ma, X., Tian, Y., Song, Y., Shi, J., Xu, J., Xiong, K., Li, J., Xu, W., Zhao, Y., Shuai, J., Chen, L., Plikus, M. V., Lengner, C. J., Ren, F., Xue, L., and Yu, Z. (2017) Msi2 Maintains Quiescent State of Hair Follicle Stem Cells by Directly Repressing the Hh Signaling Pathway. J. Invest. Dermatol. 137, 1015–1024

46. Cragle, C., and MacNicol, A. M. (2014) Musashi Protein-directed Translational Activation of Target mRNAs Is Mediated by the Poly(A) Polymerase, Germ Line Development Defective-2. J. Biol. Chem. 289, 14239–14251

47. Rutledge, C. E., Lau, H.-T., Mangan, H., Hardy, L. L., Sunnotel, O., Guo, F., MacNicol, A. M., Walsh, C. P., and Lees-Murdock, D. J. (2014) Efficient Translation of Dnmt1 Requires Cytoplasmic Polyadenylation and Musashi Binding Elements. PLOS ONE. 9, e88385

48. MacNicol, M. C., Cragle, C. E., McDaniel, F. K., Hardy, L. L., Wang, Y., Arumugam, K., Rahmatallah, Y., Glazko, G. V., Wilczynska, A., Childs, G. V., Zhou, D., and MacNicol, A. M. (2017) Evasion of regulatory phosphorylation by an alternatively spliced isoform of Musashi2. Sci. Rep. 7, 1–17

